# Phylogenetic position and mitochondrial genome evolution of ‘orphan’ eukaryotic lineages

**DOI:** 10.1101/2025.04.16.649146

**Authors:** Vasily Zlatogursky, Vittorio Boscaro, Gordon Lax, Matias Wanntorp, Nina Pohl, Fabien Burki, Patrick J. Keeling

**Affiliations:** Department of Botany, University of British Columbia, Vancouver, BC, Canada; Department of Organismal Biology, Uppsala University, Uppsala, Sweden

## Abstract

The phylogenetic tree of eukaryotes is divided into a handful of highly diverse ‘supergroups’; only a few so-called ‘orphan’ lineages branch in uncertain positions outside of these large clades. We found that the mitochondrial genome of one such lineage, the telonemids, is considerably gene-rich, a feature observed in other ‘orphans’ as well, raising the possibility that these organisms share a common history. On the contrary, our phylogenomic analyses show that ‘orphans’ with gene-rich mitochondria branch into two different positions: telonemids actually belong to the established supergroup Haptista, while provorans and meteorids form a strongly supported clade with hemimastigophorans, in a novel ancient supergroup that we dub here Promethea. Comparative genomics shows that this split reflects differences between mitochondrial gene sets. Thanks to the increased number of available representatives analyzed together, our results further simplify and illuminate the evolutionary relationships between eukaryotes.

## Introduction

Eukaryotes include familiar organisms like animals, plants, and fungi, but also dozens of protist groups that are just as old and diverse, and their evolutionary relationships – the ‘tree of eukaryotes’ – have been the subject of intense investigation. In the last decades, large-scale phylogenomic analyses have slowly but surely converged into a consensus that identifies a small number of ‘supergroups’^1-4^, far broader than traditional ‘kingdoms’, as the main blocks of eukaryotic diversity. For example, animals and fungi are just two of the subdivisions within one supergroup, Opisthokonta, while plants originated from within one of several major lineages of algae in the supergroup Archaeplastida.

While these large clusters were being defined, new microbial eukaryotes were also steadily being discovered. Many fell within the established supergroups, but a handful of so-called ‘orphan’ lineages were found to branch outside, even when using genome-wide phylogenetic analyses, and their relationships with one another and other eukaryotes remain uncertain. Subsets of four such orphans, i.e. the provorans^5^, meteorids^6^, hemimastigophorans^7^, and telonemids^8,9^, occasionally branch together in phylogenomic analyses^6,10-12^, although inconsistently. Moreover, the available mitochondrial genomes from two of them (provorans and meteorids) are remarkably gene-rich^5,6,13^. The possibility that all four of these lineages that do not neatly fit into the emerging framework of eukaryotic relationships might form a single clade with ancestrally gene-rich mitochondria is hence a reasonable hypothesis to test. Unfortunately, these groups are all underrepresented and seldom included together in the same analysis – in part due to the fact that some of them have been discovered or molecularly investigated very recently, and virtually at the same time – so there has been no specific effort to address this question.

Despite having been studied for far longer than the others, the position of telonemids is probably the most unstable and controversial, both because different molecular analyses place them in different parts of the tree and because their unique ultrastructural features do not univocally suggest a close relationship with any other organism^10,14,15^. Similarly, there are very few morphological similarities between hemimastigophorans, meteorids, and provorans, although the sister group status of the first two has been determined with high confidence^6^.

Here, we characterize the transcriptome and the mitochondrial genome of a newly described telonemid, *Microkorses curacao* gen. et sp. nov., found in the Caribbean. We observe that its mitogenome is gene-rich, superficially reminiscent of those of other ‘orphan’ groups, but in fact retaining different genes than in provorans and meteorids. Our phylogenomic analyses furthermore demonstrate that provorans, meteorids, and hemimastigophorans form a single, well-supported but previously unrecognized supergroup, established here as ‘Promethea’, while telonemids branch within an existing supergroup, Haptista.

## Results and Discussion

### A new telonemid

Telonemids are small, heterotrophic unicellular protists with two flagella and a mosaic of ultrastructural features that are either wholly unique or resemble those of various other groups^9,16^. Environmental DNA analyses suggest that they are ubiquitous in aquatic environments^17,18^, but there are currently only three formally described genera: *Telonema, Lateronema* and *Arpakorses*, with only a few species each. We here present a new genus and species of telonemids, *Microkorses curacao* gen. et sp. nov., isolated from the plankton of the Caribbean Sea around Curaçao. It was represented by nanoflagellates (5-6 µm) which were mostly sedentary under the culture conditions, but were also capable of fast spiral swimming in the water column (Fig. 1; Supplementary videos S1-3). Despite its tiny size, *M. curacao* is a voracious predator, able to efficiently clear a dense culture of bigger (7-12 µm) eukaryotic prey (the kinetoplastid, *Neobodo designis*) in a couple of weeks. *M. curacao* belongs to the TEL1 clade ^16^ with *Telonema* and *Arpakorses* (Suppl. Fig. S1), and it is the smallest telonemid described so far; other distinguishing features include two flagella of equal length and a tendency to attach to the substratum (see Supplementary Discussion for a full characterization).

**Figure 1.**
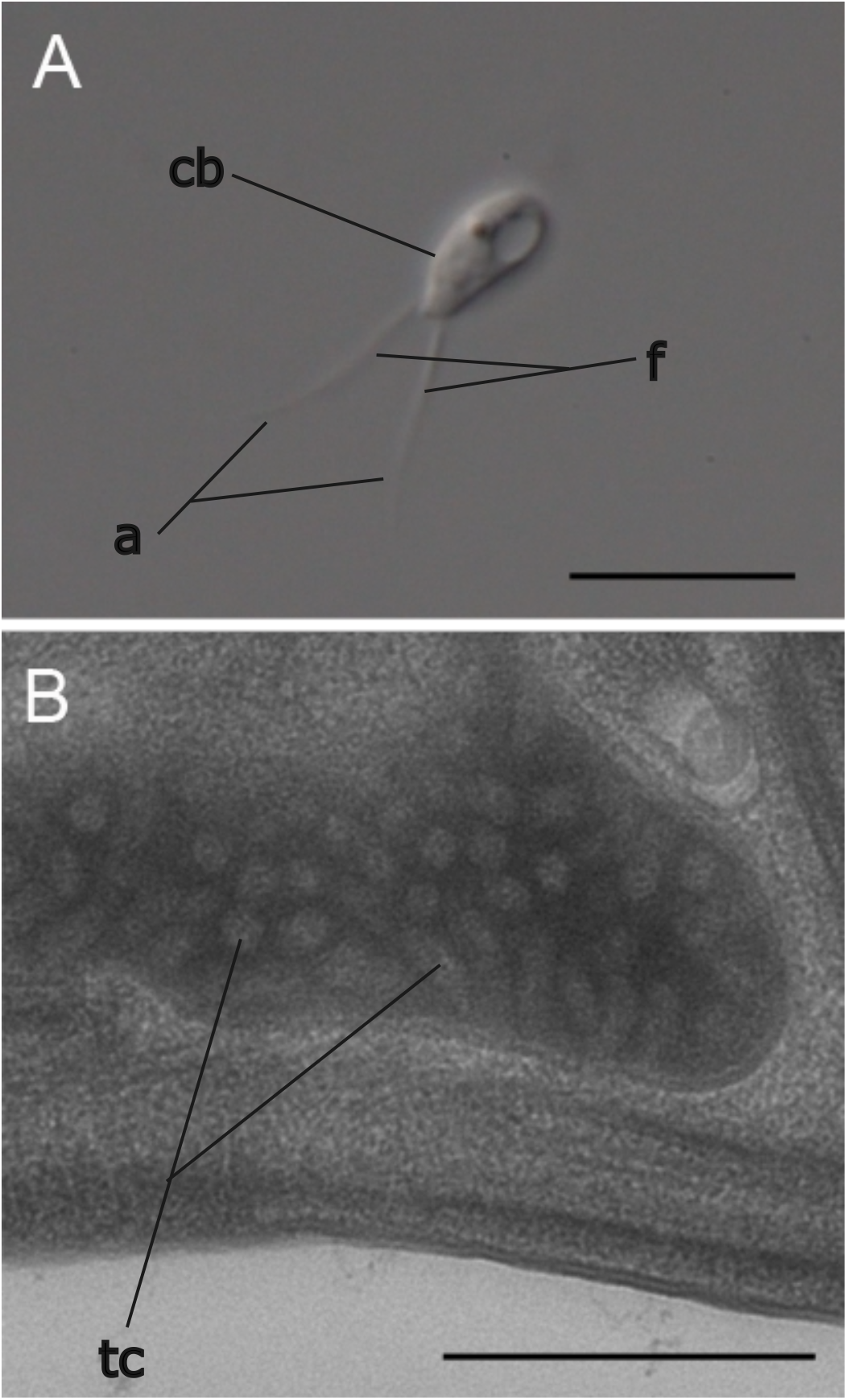
Cell morphology of *Microkorses curacao* gen. et sp. nov. A. General view of the cell. Light microscopy, differential interference contrast. The scale bar stands for 5 µm. B. The mitochondrion with tubular cristae is visible in this transmission electron microscopy cross-section. Abbreviations: a - acronemes, cb - cell body; f - flagella, tc - tubular cristae. The scale bar stands for 500 nm.

### Mitochondrial genome features of telonemids

Starting from a clonal culture strain (CU2384), we sequenced the complete mitochondrial genome of *M. curacao*, obtaining a circular-mapping sequence 42,908 bp long, with a G+C content of 26.5%, that closely matches previously obtained environmental data^19^ (Supplementary Discussion, Suppl. Fig. S2, Suppl. Tables S1-S2). The mitogenome encodes 74 genes, 45 of which protein-coding; of these, 5 are unidentified open reading frames (ORFs), and 3 are ribosomal genes (*rnl, rns, rrn5*). There are genes coding for 26 tRNAs collectively capable of recognizing all 20 amino acids, but not stop codons. The telonemid mitogenome encodes 40 mitochondrial protein-coding genes out of the 74 known to theoretically exist^19,20^, altogether making it considerably gene-rich: by comparison jakobids, the eukaryotic lineage with the most gene-rich mitochondrial genomes, have up to 66 protein-coding genes^21^. Richer genomes^6,20,22^ can only be found in some discobans, CRuMs, cryptists, and *Malawimonas jakobiformis*, as well as the new supergroup described here (see below), the prometheids (Fig. 2). The mitogenome of *M. curacao* contains some rarely retained genes coding for proteins of the large ribosomal subunit, such as *rpl10, rpl31* and *rpl32*; this feature is shared by prometheids. However, *rpl18* is missing in prometheids but present in *M. curacao*, while *rpl20* and *rpl27* are by contrast only retained in prometheids. Both lineages share some uncommon genes, such as *tatC*, but no signs of heme biosynthesis or succinate dehydrogenase (complex II) genes were found in the *M. curacao* mitogenome nor in its transcriptome, while being well represented in prometheids. This points to the retention of different sets of mitogenome-encoded proteins in prometheids and telonemids. The *M. curacao* mitogenome content is relatively similar to that of centrohelids, especially the ribosomal protein set (Fig. 2), but it is considerably more gene-rich than the mitogenome of the other major group of Haptista, the haptophytes (again, especially in ribosomal protein genes). Overall, the extended repertoire of mitochondrial genes in telonemids does not suggest any especially close relationship to meteorids or provorans (whose mitogenomes do instead resemble each other’s^6^), indicating that rich mitochondrial genomes are not a synapomorphy uniting all these ‘orphan’ groups. Rather, telonemids and prometheids have independently retained different sets of genes from the ancestral eukaryotic mitochondrion (a less likely alternatively is that either or both re-acquired some of these genes by horizontal gene transfer, which however happens at very low rates in the mitochondria of most lineages).

**Figure 2.**
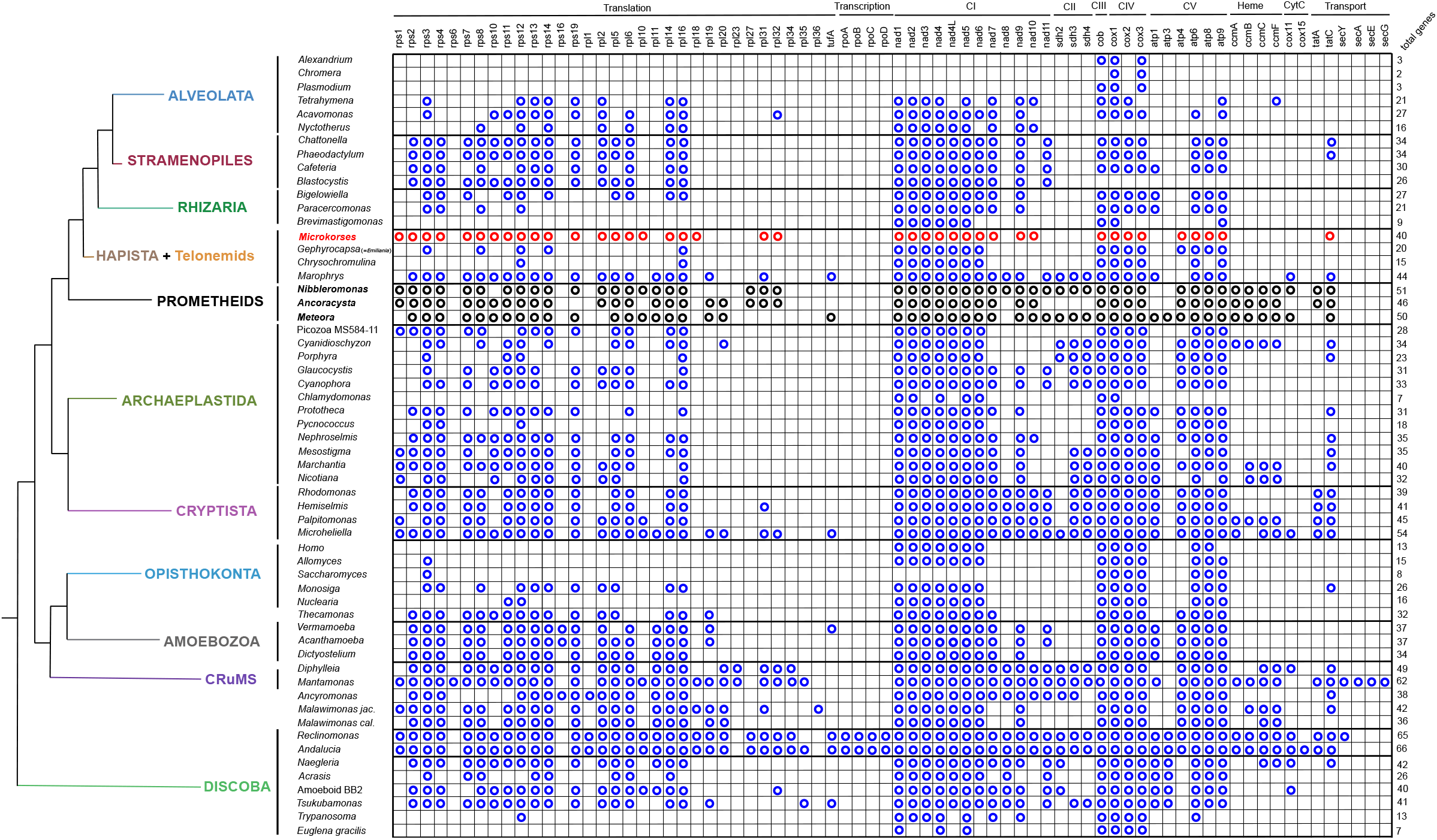
Eukaryotic mitogenomes coding capacity. Comparison of presence (circle) / absence (no circle) of protein-coding genes in mitochondrial genomes of eukaryotes (updated and corrected when necessary from previous sources^6,20,22^). The *Microkorses* branch is shown in red, prometheids are in bold and in black. The depicted cladogram is based on the tree shown in Fig. 3. Abbreviations: CI-CV - Respiratory complexes I- V; CytC - Cytochrome C.

### Placement of telonemids and other ‘orphan’ groups in the tree of eukaryotes

Telonemids have been poorly represented in phylogenomic datasets until recently^15^, while other ‘orphan’ lineages, specifically provorans, meteorids, and hemimastigophorans, were molecularly characterized in rapid succession^5-7^ and hence never all included with a representative number of species in the same phylogenomic tree. To analyse these orphan groups together in the context of eukaryote evolution, we built two alignment matrices using 278 protein-coding genes: one including 89 taxa, chosen to optimize both completeness and taxonomic breadth, and a much larger one with 433 taxa, vastly increasing the number of representatives from all eukaryotic groups. The 89-taxa Maximum-Likelihood (ML) tree was inferred using two complex models (LG+C60+G4 and LG+UDM64+G4) which produced identical topologies (Fig. 3). Bayesian inference did not converge even after 1,500 iterations (maxdiff: 1), although notably all four chains provided full support for the position of telonemids recovered by ML methods. We inferred instead a tree using the hybrid CAT-PMSF method^23^, which coincided with the ML topologies in all but three nodes (Fig. 3). The 433-taxa ML tree was inferred with a less parameter-rich model (LG+C20+G4) and produced a similar result (Suppl. Fig. S3). In particular, all analyses showed telonemids branching with Haptista, although the exact relationship between telonemids and the two major haptist subgroups (haptophyte algae and the heterotrophic centrohelids) could not be determined with confidence. Provorans, meteorids, and hemimastigophorans always formed a single clade, here dubbed Promethea from the initial letters of their names and after the mythological Greek titan, as a nod to the ancient origin of their last common ancestor. Prometheids and haptists in turn branched within a very diverse clade also containing SAR, Archaeplastida, and Cryptista (the latter two always being sister groups in all our analyses), which has been reported by many phylogenomic analyses6,10,15,24.

**Figure 3.**
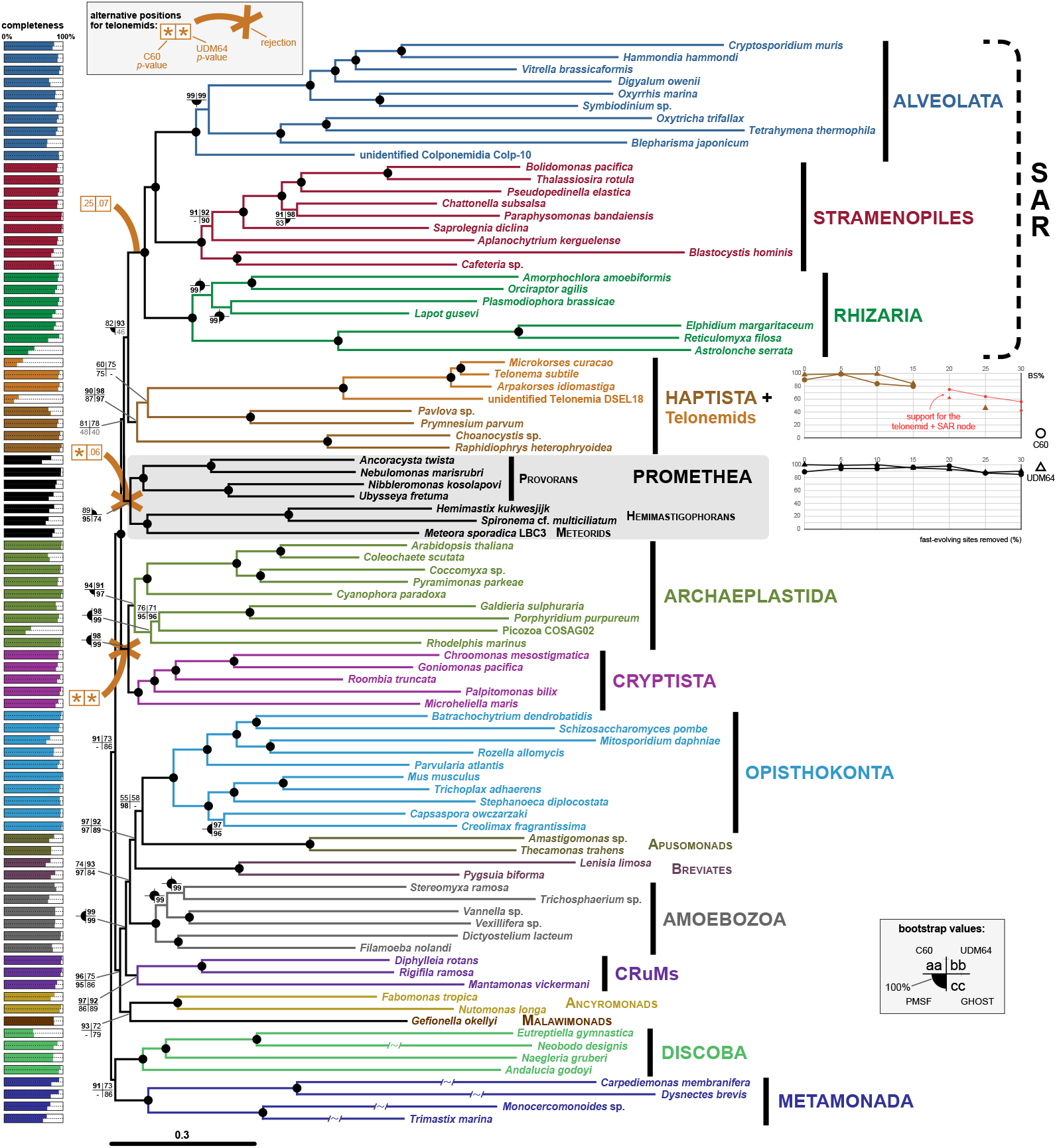
The position of telonemids and prometheids in the tree of eukaryotes. Multigene Maximum-Likelihood phylogenetic tree of eukaryotes, showing that telonemids belong to Haptista while hemimastigophorans, provorans, and meteorids form a single clade, Promethea. The character matrix for 89 representative eukaryotes was built with 278 protein-coding genes and 73,081 sites. The shown tree was inferred with the LG+C60+G4 model. Other bootstrap values for each node were calculated with the LG+UDM64+G4 and LG+C20+H4 models as well as the CAT-PMSF method, as shown in the inset. 100% bootstrap scores are shown as black circles ; missing values indicate that the node was not recovered in the corresponding analysis. For space reasons, the four terminal branches in Metamonada and one terminal branch in Discoba were shortened by 50%. The completeness of each taxon’s genome or transcriptome is shown on the left, as a percentage of represented genes (top bars) and non-gap sites (bottom bars). Alternative positions for telonemids assessed with AU tests are shown (asterisks in the two boxes correspond to a rejection with the LG+C60+G4 model, on the left, and the LG+UDM64+G4, on the right). Graphs next to the Haptista and prometheid clades show bootstrap value changes when fast-evolving sites are removed (the absence of a datapoint denotes that the node was not recovered with the corresponding matrix); support for the alternative position of telonemids obtained with smaller matrices is also shown, in red. Note that the position of the root was chosen arbitrarily.

We then focussed on testing the stability of the two most novel findings: the node supporting the monophyly of telonemids and haptists, and the node supporting the monophyly of prometheids. To determine the impact of substitution rate heterogeneity across sites, we repeated the 89-taxa dataset ML analyses on matrices with incremental removal of fast-evolving sites; the monophyly of prometheids was always recovered with very high support; support for the monophyly of telonemids and haptists increased when the most extreme fast-evolving sites were removed, but eroded or disappeared once 20% or more of the information was lost (Fig. 3). In these reduced matrices, telonemids branched as the sister group of SAR, but with low bootstrap support, further reduced as the information decreases (down to 43% after removing 30% of the sites). The prometheid and haptists+telonemid clades were also recovered, with similar or higher support values, in analyses that only included the taxa with the highest data completeness (Suppl. Fig. S4), and when removing faster-evolving taxa (Suppl. Fig. S5). The effect of lineage-specific differences in substitution rates was further addressed using the GHOST model^25^ on the 89-taxa dataset, which confirmed the two aforementioned nodes as well as most of the topology recovered under the other approaches (Fig. 3). Telonemids were previously found to branch in several different parts of the tree of eukaryotes, including as sister to SAR in the ‘TSAR’ hypothesis^15^. To test this and other proposed or plausible placements, we built ML trees where telonemids were constrained to cluster with either SAR, prometheids, or in a clade with Archaeplastida and Cryptista. The ‘TSAR’ scenario was not rejected by Approximately Unbiased tests (*p*-values: 0.25 and 0.07 with the LG+C60+G4 and LG+UDM64+G4 models, respectively), while the clustering of telonemids with prometheids was rejected by the C60 model (*p*-value: 0.06 with the LG+UDM64+G4 model); a relationship with Archaeplastida and Cryptista was rejected by both models (Fig. 3).

The monophyly of prometheids is not a complete surprise. While no tree encompassing their whole diversity was reconstructed before, meteorids and hemimastigophorans have been shown to branch together, and on occasion to be related with the provoran *Ancoracysta*^6,12^, despite their completely different morphologies and modes of locomotion. Meteorids are non-flagellated, gliding protists with two characteristic lateral ‘arms’ swinging back and forth, while hemimastigophorans bear multiple flagella (or ‘cilia’) arranged in two lateral rows (kineties) – and yet, while sister groups, they are approximately as divergent as the two superficially similar lineages of provorans (nibblerids and nebulids) are from each other (Fig. 3 and Suppl. Fig. S3). Provorans are biflagellated protists that voraciously feed on other eukaryotes, using extruding organelles (which have been suggested to have homologs in hemimastigophorans) and small but strong sub-cellular ‘jaws’^26^. While more data is needed, especially from the mitochondria of hemimastigophorans, it is reasonable to assume that the prometheid ancestor had a gene-rich mitogenome, as is still the case in its extant descendants.

The placement of telonemids is much more controversial. Interestingly, the first phylogenomic analysis of telonemids had them branch with centrohelids^14^, but this association was not observed in later trees, which recovered various topologies, of which ‘TSAR’ is only one^10,11,12^. Multiple authors have remarked that different morphological features of telonemids could point to different relatives^15,16^ or to none, as is the case for their distinctive cytoskeleton^27^. Telonemid sequences are not especially fast-evolving, so the reason behind their erratic behaviour in phylogenomic trees must lie elsewhere, for example in their sparse sampling and the lack of close relatives.

### Promethea Zlatogursky, Boscaro, and Keeling clad. nov

Diagnosis: The most inclusive clade containing *Ancoracysta twista* Janoušcovec, Tikhonenkov, Burki, Howe, Rohwer, Mylnikov & Keeling 2017, *Hemimastix kukwesjijk* Eglit & Simpson 2018, and *Meteora sporadica* Hausmann, Weitere, Wolf & Arndt 2002, but not *Telonema subtile* Griessman 1913, *Paramecium aurelia* Ehrenberg 1838, or *Arabidopsis thaliana* (L.) Heynhold 1842.

The name refers to the initial letters of the three known lineages at the time of establishment (PROvorans, METeorids, and HEmimastigophorans) and to the mythological titan Prometheus, as a nod to the fact that the last common ancestor of the clade is likely to be extremely ancient. Reference tree: Fig. 3 in this publication.

## Conclusions

While the catalogue of eukaryotic diversity keeps growing, we show here that several orphan groups, until now in uncertain relationships with each other and with known supergroups, can be placed with confidence in the tree when analysed together. A few years ago, three other orphan lineages that lacked unifying morphological features were recognized based on molecular phylogenies to form a monophyletic group, CRuMs^28^. The prometheids are even more varied; for example, each of them represents one of three main forms of locomotion among protists: ciliary swimming, flagellar swimming, and gliding. Yet our analyses show that they form a strongly supported single clade. Their last common ancestor existed so long ago that it is not surprising its descendants don’t resemble each other (not to mention, more prometheid diversity will undoubtedly be uncovered in the future) - and yet, they all share a similarly gene-rich mitochondrial genome. We also show that another elusive group, the telonemids, also has a gene-rich mitogenome but is not related to the prometheids, according to both comparative genomics and phylogenomics. The telonemids’ most likely placement is within the well-established supergroup, Haptista, based on the inclusion of many representatives for each group of eukaryotes, multiple models, and multiple taxon selections, including one particularly extensive phylogenomic tree (Suppl. Fig. S3). This conclusion is nevertheless still not definitive, since one proposed alternative, the so-called ‘TSAR’ hypothesis, could not be rejected. Removing these four lineages from the category of ‘orphans’ further solidifies a single and comprehensive tree that will be a useful tool as we interpret the incredible diversity of eukaryotic cells.

## Limitations of the study

Two particular courses of action would strengthen our conclusions. First, the oft-cited alternative for the position of telonemids, i.e. their being the sister group of SAR (‘TSAR’ hypothesis), was not rejected by statistical tests. It should be noted that most of our trees (although not all, and with varying support) depict Haptista (including telonemids) as the sister group of SAR – this might mean that in the TSAR topology it was not the telonemids’ position, but rather that of other haptists, that was incorrect. If true, this in turn might require obtaining more data from underrepresented haptists (for example, centrohelids) in order to stabilize the topology. Secondly, there is currently no data on the mitochondrial genomes of hemimastigophorans. Until these become available, the synapomorphy status of the gene-rich mitogenome for Promethea cannot be confirmed.

## Supporting information

Supplementary discussion

## Resource availability

### Lead contact

Requests for further information and resources can be directed to the lead contact, Vasily Zlatogursky.

### Materials availability

This study did not generate new unique reagents.

### Data and code availability

- The *Microkorses curacao* mitochondrial genome sequence and transcriptomic raw reads have been deposited on GenBank as [accession number pending] and are publicly available as of the date of publication.
- Alignments used in this study have been deposited on the Borealis data repository [link pending].
- This paper does not report original code.
- Any additional information required to reanalyze the data reported in this paper is available from the lead contact upon request.

## Acknowledgements

Authors are thankful to Samuel Livingston for assistance with electron microscopy, Mahara Mtawali for collecting the sample, and Adarsh Hurdeal for helping with imaging. This work was supported by grants from the Natural Sciences and Engineering Research Council of Canada (NSERC 2019-03994) and the Gordon and Betty Moore Foundation (https://doi.org/10.37807/GBMF9201) to PJK, and by grants from the European Research Council (ERC consolidator grant 101044505), the Swedish Research Council VR (2021-04055), and Science for Life Laboratory to FB.

## Author contributions

V.Z., V.B., and P.J.K. designed the project. V.Z. isolated and cultured the organisms, performed microscopy and molecular biology experiments, and analyzed mitochondrial data. V.B. performed phylogenetic analyses, with assistance from G.L. M.W., N.P., and F.B. curated the starting PhyloFisher database. V.Z., V.B., and P.J.K. wrote the original draft of the paper. All authors contributed to the final manuscript.

## STAR methods

Detailed methods are provided in the online version of this paper and include the following:

*<u>Table</u> for STAR*

### Isolation, culturing, and morphology of *Microkorses curacao* gen. n., sp. n

*Microkorses curacao* strain CU2384 was isolated from a plankton-tow collected water sample taken from the Caribbean Sea at the water factory dive site in Willemstad, Curaçao on October 10^th^, 2023 (latitude/longitude coordinates: 12.108993807580502, -68.95353029793979). The salinity measured in the sample was 36‰. The sample was transported to the laboratory and inoculated in a 90 mm plastic Petri dish. The dish was incubated at room temperature for two weeks with addition of 0.025% wheat grass extract to promote bacterial growth. The incubated dish was examined using a Zeiss AxioVert A1 inverted microscope, equipped with phase contrast (PC). The living cells of *Microkorses* were manually picked using tapered glass Pasteur pipettes and inoculated individually into wells of a plastic 96-well plate, containing 200 μl of actively growing *Neobodo designis* culture in 0.025% wheat grass extract in 30 ppt artificial sea water. One well, where a clone of cells was noted to be successfully growing, was fully transferred to a Petri dish with *N. designis*, and the obtained clonal culture was further maintained by bi-weekly reinoculation of 1 ml of old culture onto a fresh dish with the prey. The culture was maintained at 15 °C.

Light microscopy was performed in plastic Petri dishes using a Zeiss AxioVert A1 with PC or on temporary object slide preparations on the same microscope, but also on a Zeiss Axioplan 4 upright microscope, equipped with differential interference contrast (DIC) optics. The photographs and videos were captured with a Sony alpha-7R Mark 3 camera. For transmission electron microscopy the cells were fixed using high-pressure freezing (HPF). First, the cells were concentrated by centrifugation at 560 *g* for 10 minutes, the supernatant was then discarded and 1.4 μl of the concentrated cells was applied to brass planchets and immediately frozen using a Leica HPM100 HPF system. The frozen planchets were transferred to cryovials containing 2% OsO_4_ and 0.1% uranyl acetate (UA) in anhydrous acetone and processed in an automatic freeze-substitution system (Leica AFS2) according to the following schedule: 96 hours at -90 °C, heated to -50 °C over 12 hours, left at -50 °C for 8 hours, heated to -20 °C over 12 hours, left at -20 °C for 8 hours, then heated to 22 °C for 12 hours. Samples were rinsed 3 times in anhydrous acetone and then infiltrated in an ascending graded series of Spurr’s resin in acetone (20%, 40%, 60%, 80%, then three infiltrations of 100%) for a minimum of 3 hours per step. Samples were polymerized at 60 °C for 48 hours. Sections were cut from Spurr’s embedded blocks using a Leica UC7 Ultramicrotome and collected on formvar-coated copper grids, post-stained with 2% aqueous UA and lead citrate and imaged using a Tecnai Spirit TEM operating at 80 kV with a DVC1500M side-mounted camera. The size of the microscopic structures was measured on micrographs using ImageJ^29^ version 1.54f. Size measurements were performed on 185 individual cells.

### Transcriptome and mitogenome sequencing and assembly

In order to obtain single-cell transcriptomes, the cells were manually picked and frozen in 0.2 mL plastic tubes containing 2 μl of the lysis buffer. Cells were collected during the later stages of culture growth, once no prey had been detected under the microscope for several days; the organisms were washed multiple times in fresh medium to further reduce any contamination. Five cells were combined in a tube in order to increase cDNA yield. The downstream processing was performed following the Smart-seq2 protocol^30,31^ using 24 PCR cycles. For genomic sequencing, the cells were concentrated at 2,700 *g* for 10 minutes, then the total genomic DNA was extracted using the DNeasy Plant Pro kit (Qiagen). Both cDNA and genomic DNA were then sequenced with Illumina. Illumina (Nextera XT) libraries were prepared and sequenced on the Illumina NovaSeq platform at the Sequencing and Bioinformatics Consortium at the University of British Columbia. Raw reads (available in the Sequence Read Archive; accession number pending) were error-corrected using Rcorrector^32^ version 1.0.4 and trimmed using Trimmomatic^33^ version 0.39 with the options LEADING:5 TRAILING:5 SLIDINGWINDOW:5:16 MINLEN:60, specifying Nextera adapters, Illumina-specific primers and smartseq2 primers for removal. The transcriptome was assembled with SPAdes^34^ version 3.15.1 using reverse and unpaired Illumina reads, and the --rna flag. The obtained transcriptome contained 1,141 contigs, had N50 of 1,415 bp, and contained 5.1% of conserved eukaryotic genes according to BUSCO^35^ version 5.7.1 and 1,217 predicted peptides. Protein-coding sequences were predicted from the assemblies using Transdecoder^36^ version 5.5.0, then processed through the PhyloFisher pipeline (see below); each orthogroup was manually inspected in single-gene trees, removing paralogs that did not cluster with previously sequenced telonemids (mostly originating from the prey organism, *Neobodo designis*, which clusters in a completely different part of the tree, with other representatives of its genus).

Assembly of the *M. curacao* mitochondrial genome was performed on SPAdes as described above for the transcriptome, using the --meta flag. The assembly was blasted using barrnap-obtained mitochondrial SSU sequence as a query, which resulted in the recovery of a single circular contig. Annotation was performed using the MFannot webserver^37^, and the obtained .sqn file was converted to GenBank format using asn2gb (https://ftp.ncbi.nlm.nih.gov/asn1-converters/by_program/asn2gb/). In parallel, the same fasta file was annotated with PROKKA^38^ version 1.14.6 using the --kingdom Mitochondria flag. The MFannot file was manually corrected based on the PROKKA output wherever it was better performing. The genome map was generated using OGDRAW^39^ version 1.3.1.

### Phylogenetic analyses

SSU rRNA gene sequences were extracted from the obtained transcriptome using barrnap version 0.9 (https://github.com/tseemann/barrnap). Other telonemid sequences were downloaded from the PR2 database^40^ and manually supplemented with some additional sequences individually downloaded from GenBank. The resulting fasta file was clustered at 99% to avoid redundancy using usearch^41^ version 12, aligned using mafft^42^ version 7.48 and trimmed using trimAl^43^ version 1.4.r22 with a gap threshold of 0.3 and similarity threshold of 0.001. The obtained file was used for Maximum-Likelihood (ML) analysis using RAxML-NG^44^ version 1.1.0 under the GTRCAT model and 1,000 non-parametric bootstrap replicates. Visualisation of this and subsequent trees was done with iTOL^45^ version 6.9.1, FigTree version 1.4.4 (https://tree.bio.ed.ac.uk/software/figtree/), InkScape version 1.2, and Adobe Illustrator version 27.5.

The newly obtained telonemid transcriptome and other genomic and transcriptomic data recently made available (mostly collected from EukProt^46^, Supplementary Table S3) were added to a previous dataset^4^ formatted for PhyloFisher^10^. 320 single-gene trees of 1,129 taxa were inferred with the PhyloFisher sgt_constructor.py pipeline, then curated manually to remove paralogs and suspected contaminants. At the end of this preliminary step, genes present in less than 50% of the taxa, as well as a few genes whose tree topology was too ambiguous or too complex (for example, due to large duplication events) to confidently curate, were removed, leaving 278 genes. After removing bacterial and archaeal data, taxa with less than 10% site coverage, the notoriously fast-evolving Ascetosporea^47^, and nucleomorph data, 793 taxa remained. Preliminary phylogenomic trees were built on this comprehensive dataset. 17 very long-branching taxa were further removed; many others were pruned due to redundancy (multiple taxa from the same genus or species, and/or closely related to each other; in each case, the most complete representative was chosen). This left the 433 taxa used in the large dataset analyzed in the paper; 89 of these, chosen to maintain a wide taxonomic breadth while maximizing completeness (measured as both gene- and site-percentage representation in the supermatrix) whenever possible, constituted the smaller dataset. The alignment matrix in both datasets included 278 genes and 73,081 sites.

All multi-gene ML inferences were performed on IQ-TREE^48^ version 2.2.0. The 433-taxa dataset was used to infer a ML tree with the LG+C20+G4 model, running 1,000 ultrafast bootstraps for node support. ML trees were generated for the 89-taxa dataset using the more complex LG+C60+G4 and LG+UDM64+G4 models, running 1,000 ultrafast bootstraps for node support. To assess the impact of sites with higher substitution rates, the 5% fastest-evolving sites were removed from the original 89-taxa alignment using the fast_site_removal.py script, in incremental steps up to 30%. Ultrafast bootstrapped ML trees were run on each submatrix, extracting the boostrap values of nodes of interest. Additional ML inferences based on both the LG+C60+G4 and the LG+UDM64+G4 models were run on datasets including only taxa with 75% or more gene coverage (81 out of the original 89) and only taxa with shorter branches (78 out of 89). In unrooted trees (as all ours are, given the lack of consensus on the position of the root of eukaryotes) there is no single way to define the total branch length of a taxon, so the choice of which taxa to remove was based on the unevenness branch lengths between related taxa; this included all metamonads, *Neobodo designis*, all retarians (i.e. *Astrolonche serrata, Reticulomyxa filosa*, and *Elphidium margaritaceum*), *Blastocystis hominis, Tetrahymena termophila*, and *Cryptosporidium muris*.

A Bayesian Inference analysis was attempted under the CAT-GTR model on PhyloBayes^49^ version 1.8c, but runs have not converged after ∼1,500 generations. In its stead, we employed the Bayesian / ML hybrid CAT-PMSF method^23^ on the 89-, 81-, and 78-taxa datasets, using ML reference trees obtained with the LG+F+G4 model to calculate site frequencies and exchangeability rates on PhyloBayes (model: CAT-GTR); these were then used, combined with a FreeRate model with 4 categories, to infer a final ML tree on IQ-TREE, running 1,000 ultrafast bootstraps. The computationally intensive GHOST model^25^, which was designed to account for branch-dependent heterotachy, was also run on the 89-taxa dataset (−m LG+C20+H4 command, running 1,000 ultrafast bootstraps).

Constrained trees were built to test alternative topologies on the 89-taxa dataset. In each case, a few nodes were fixed before running a full phylogenetic analysis with the LG+C60+G4 model to obtain the best topology. Specifically, the three constrained trees were run with the following nodes set: (i) telonemids (monophyletic) as sister group of SAR (monophyletic); (ii) telonemids (monophyletic) as sister group of prometheids (monophyletic); (iii) telonemids (monophyletic), Archaeplastida (monophyletic), and Cryptista (monophyletic) forming a single clade (but in no pre-defined relationship with each other). Notably, the constrained position of telonemids was the only difference between constrained trees and the unconstrained ML tree. Approximately Unbiased (AU) tests were then run in IQ-TREE (using 10,000 RELL replicates) to determine which of these three alternative topologies could be significantly rejected, according to both the LG+C60+G4 model and the LG+UDM64+G4 model.

## Supplemental information

### Suppl. Discussion

Taxonomic diagnosis and further considerations on mitochondrial gene features.

**Suppl. Fig. S1**. Maximum-Likelihood tree based on SSU rRNA gene sequences, inferred using 1,736 nucleotide positions from 120 telonemids and 10 diverse eukaryotes as an outgroup, under the GTRCAT model, with 1,000 non-parametric bootstrap replicates. The new *Microkorses curacao* species is shown in red. The SSU sequences corresponding to environmental mitogenomes are shown in blue. Morphologically characterized taxa are shown in green. Clade names are based on Fig. 1 in Tikhonenkov et al.^16^; arrows and vertical lines denote clades.

**Suppl. Fig. S2**. The map of the *Microkorses curacao* mitochondrial genome with genes colour coded as to their function, generated using OGDRAW. Genes on the outside of the circle are transcribed in the clockwise direction.

**Suppl. Fig. S3**. Maximum-Likelihood phylogenetic tree of 433 representative eukaryotes. The character matrix included 278 protein-coding genes and 73,081 sites, and the tree was inferred under the LG+C20+G4 model, running 1,000 ultrafast bootstraps for node support; 100% bootstrap scores are shown as black circles. The position of the root was chosen arbitrarily. All the main features of the 89-taxa tree are confirmed in this larger tree, including the branching of telonemids with Haptista, the monophyly of prometheids, diaphoretickes, and Archaeplastida+Cryptista.

**Suppl. Fig. S4**. Maximum-Likelihood phylogenetic tree of a subset of 81 taxa from Fig. 3 with high (75% or more) gene coverage. The shown t ree was inferred with the LG+C60+G4 model. Other bootstrap values for each node were calculated with the LG+UDM64+G4 model and the CAT-PMSF method, as shown in the inset. 100% bootstrap scores are shown as black circles; missing values (hyphens) indicate that the node was not recovered in the corresponding analysis. The completeness of each taxon’s genome or transcriptome is shown on the left, as a percentage of represented genes (top bars) and non-gap sites (bottom bars). The position of the root was chosen arbitrarily. The topology corresponds to the topology in the main analyses.

**Suppl. Fig. S5**. Maximum-Likelihood phylogenetic tree of a subset of 78 taxa from Fig. 3, after removing 11 taxa with longer branches. The shown t ree was inferred with the LG+C60+G4 model. Other bootstrap values for each node were calculated with the LG+UDM64+G4 model and the CAT-PMSF method, as shown in the inset. 100% bootstrap scores are shown as black circles ; missing values (hyphens) indicate that the node was not recovered in the corresponding analysis. The completeness of each taxon’s genome or transcriptome is shown on the left, as a percentage of represented genes (top bars) and non-gap sites (bottom bars). The position of the root was chosen arbitrarily. The topology corresponds to the topology in the main analyses.

**Suppl. Fig. S6**. The comparison of the gene content and synteny in *Microkorses curacao* and three previously obtained SAG-based mitogenomes. Genome maps are shown as linear for clarity. The vertical lines link corresponding genes. Maps were generated using OGDRAW. The annotation is colour-coded as in Fig. S2.

**Suppl. Table S1**. Comparison of the tRNA gene content in *M. curacao* and three previously obtained SAG-based mitogenomes.

**Suppl. Table S2**. A table supplementing Fig. 2 and showing mitogenome completeness across eukaryotes.

**Suppl. Table S3**. Identity and source of the phylogenomic data added to the starting dataset^4^.

**Suppl. Video S1**. *Microkorses curacao* gen. et sp. nov. imaged using an inverted Zeiss Axiovert A1 microscope (40x and 63x objectives) with phase contrast on temporary object slide with coverslip (same slide as in Suppl. video S3).

**Suppl. Video S2**. *Microkorses curacao* gen. et sp. nov. imaged using an inverted Zeiss Axiovert A1 microscope (20x and 40x objectives) with phase contrast in a Petri dish.

**Suppl. Video S3**. *Microkorses curacao* gen. et sp. nov. imaged using an upright Zeiss Axioplan 4 microscope (100x objective) with differential interference contrast on a temporary object slide with coverslip (same slide as in Suppl. Video S1).

## References

1. Brown, M.W., Heiss, A.A., Kamikawa, R., Inagaki, Y., Yabuki, A., Tice, A. K., Shiratori, T., Ishida, K.-I., Hashimoto, T., Simpson, A.G.B., and Roger, A.J. (2018). Phylogenomics places orphan protistan lineages in a novel eukaryotic super-group. Genome Biol. Evol. 10, 427–433. 10.1093/gbe/evy014.

2. Keeling, P.J., and Burki, F. (2019). Progress towards the Tree of Eukaryotes. Curr. Biol. 29, R808–R817. 10.1016/j.cub.2019.07.031.

3. Strassert, J.F.H., Irisarri, I., Williams, T.A., and Burki, F. (2021). A molecular timescale for eukaryote evolution with implications for the origin of red algal-derived plastids. Nat. Comm. 12, 1879. 10.1038/s41467-021-22044-z.

4. Schön, M.E., Zlatogursky, V.V., Singh, R.P., Poirier, C., Wilken, S., Mathur, V., Strassert, J.F.H., Pinhassi, J., Worden, A.Z., Keeling, P.J., Ettema, T.J.G., Wideman J.G., and Burki, F. (2021). Single cell genomics reveals plastid-lacking Picozoa are close relatives of red algae. Nat. Comm. 12, 6651. 10.1038/s41467-021-26918-0.

5. Tikhonenkov, D.V., Mikhailov, K.V., Gawryluk, R.M.R., Belyaev, A.O., Mathur, V., Karpov, S.A., Zagumyonnyi, D.G., Borodina, A.S., Prokina, K.I., Mylnikov, A.P., Aleoshin, V.V., and Keeling, P.J. (2022). Microbial predators form a new supergroup of eukaryotes. Nature 612, 714–719. 10.1038/s41586-022-05511-5.

6. Eglit, Y., Shiratori, T., Jerlström-Hultqvist, J., Williamson, K., Roger, A.J., Ishida, K.-I., and Simpson, A.G.B. (2024). Meteora sporadica, a protist with incredible cell architecture, is related to Hemimastigophora. Curr. Biol. 34, 451–459. 10.1016/j.cub.2023.12.032.

7. Lax, G., Eglit, Y., Eme, L., Bertrand, E.M., Roger, A.J., and Simpson, A.G.B. (2018). Hemimastigophora is a novel supra-kingdom-level lineage of eukaryotes. Nature 564, 410–414. 10.1038/s41586-018-0708-8.

8. Griessmann, K. (1913). Über marine Flagellaten. Arch. Protistenk. 32, 1–78.

9. Shalchian-Tabrizi, K., Eikrem, W., Klaveness, D., Vaulot, D., Minge, M.A., Le Gall, F., Romari, K., Throndsen, J., Botnen, A., Massana, R., Thomsen, H.A., and Jakobsen, K.S. (2006). Telonemia, a new protist phylum with affinity to chromist lineages. Proc. Biol. Sci. 273, 1833–1842. 10.1098/rspb.2006.3515.

10. Tice, A.K., Žihala, D., Pánek, T., Jones, R.E., Salomaki, E.D., Nenarokov, S., Burki, F., Eliáš, M., Eme, L., Roger, A.J., Rokas, A., Shen, X.-X., Strassert, J.F.H., Kolísko, M., and Brown, M.W. (2021). PhyloFisher: A phylogenomic package for resolving eukaryotic relationships. PLoS Biol. 19, e3001365. 10.1371/journal.pbio.3001365.

11. Williamson, K., Eme, L., Baños, H., McCarthy, C.G.P., Susko, E., Kamikawa, R., Orr, R.J.S., Muñoz-Gòmez, S.A., Minh, B.Q., Simpson, A.G.B., and Roger, A.J. (2025). A robustly rooted tree of eukaryotes reveals their excavate ancestry. Nature 640, 974–981. 10.1038/s41586-025-08709-5.

12. Torruella, G., Galindo, L. J., Moreira, D., and López-García, P. (2025). Phylogenomics of neglected flagellated protists supports a revised eukaryotic tree of life. Curr. Biol. 35, 198–207. 10.1016/j.cub.2024.10.075.

13. Janouškovec, J., Tikhonenkov, D.V., Burki, F., Howe, A.T., Rohwer, F.L., Mylnikov, A.P., and Keeling, P.J. (2017) A new lineage of eukaryotes illuminates early mitochondrial genome reduction. Curr. Biol. 27, 3717–3724. 10.1016/j.cub.2017.10.051.

14. Burki, F., Inagaki, I., Bråte, J., Archibald, J.M., Keeling, P.J., Cavalier-Smith, T., Sakaguchi, M., Hashimoto, T., Horak, A., Kumar, S., Klaveness, D., Jokobsen, K.S., Pawlowski, J., and Shalchian-Tabrizi, K. (2009). Large-scale phylogenomic analyses reveal that two enigmatic protist lineages, Telonemia and Centroheliozoa, are related to photosynthetic chromalveolates. Genome Biol. Evol. 1, 231–238. 10.1093/gbe/evp022.

15. Strassert, J.F.H., Jamy, M., Mylnikov, A.P., Tikhonenkov, D.V., and Burki, F. (2019). New phylogenomic analysis of the enigmatic phylum Telonemia further resolves the eukaryote tree of life. Mol. Biol. Evol. 36, 757–765. 10.1093/molbev/msz012.

16. Tikhonenkov, D.V., Jamy, M., Borodina, A.S., Belyaev, A.O., Zagumyonnyi, D.G., Prokina, K.I., Mylnikov, A.P., Burki, F., and Karpov, S.A. (2022). On the origin of TSAR: morphology, diversity and phylogeny of Telonemia. Open Biol. 12, 210325. 10.1098/rsob.210325.

17. Bråte, J., Klaveness, D., Rygh, T., Jakobsen, K.S., and Shalchian-Tabrizi, K. (2010). Telonemia-specific environmental 18S rDNA PCR reveals unknown diversity and multiple marine-freshwater colonizations. BMC Microbiol. 10, 168. 10.1186/1471-2180-10-168.

18. Boukheloua, R., Mukherjee, I., Park, H., Šimek, K.,Kasalický, V., Ngochera, M., Hans-Peter Grossart, H.-P., Picazo-Mozo, A., Camacho, A., Cabello-Yeves, P.J., et al. (2024). Global freshwater distribution of Telonemia protists. ISME J. 18, wrae177. 10.1093/ismejo/wrae177.

19. Wideman, J.G., Monier, A., Rodríguez-Martínez, R., Leonard, G., Cook, E., Poirier, C., Maguire, F., Milner, D.S., Irwin, N.A.T., Moore, K., et al. (2020). Unexpected mitochondrial genome diversity revealed by targeted single-cell genomics of heterotrophic flagellated protists. Nat. Microbiol. 5, 154–165. 10.1038/s41564-019-0605-4.

20. Moreira, D., Blaz, J., Kim, E., and Eme, L. (2024). A gene-rich mitochondrion with a unique ancestral protein transport system. Curr. Biol. 34, 3812–3819. 10.1016/j.cub.2024.07.017.

21. Gray, M.W., Burger, G., Derelle, R., Klimeš, V., Leger, M.M., Sarrasin, M., Vlček Č Roger, A.J., Eliáš, M., and Lang, B.F. (2020). The draft nuclear genome sequence and predicted mitochondrial proteome of Andalucia godoyi, a protist with the most gene-rich and bacteria-like mitochondrial genome. BMC Biol. 18, 22. 10.1186/s12915-020-0741-6.

22. Kamikawa, R., Shiratori, T., Ishida, K.-I., Miyashita, H., and Roger, A.J. (2016). Group II intron-mediated trans-splicing in the gene-rich mitochondrial genome of an enigmatic eukaryote, Diphylleia rotans. Genome Biol. Evol. 8, 458–466. 10.1093/gbe/evw011.

23. Szánthó, L.L., Lartillot, N., Szöllösi, G.J., and Schrempf, D. (2023). Compositionally constrained sites drive long-branch attraction. Syst. Biol. 72, 767–780. 10.1093/sysbio/syad013.

24. Burki, F., Shalchian-Tabrizi, K., and Pawlowski, J. (2008). Phylogenomics reveals a new ‘megagroup’ including most photosynthetic eukaryotes. Biol. Lett. 4, 366–369. 10.1098/rsbl.2008.0224.

25. Crotty, S.M., Minh, B.Q., Bean, N.G., Holland, B.R., Tuke, J., Jermiin, L.S., and Von Haeseler, A. (2020). GHOST: recovering historical signal from heterotachously evolved sequence alignments. Syst. Biol. 69, 249–264. 10.1093/sysbio/syz051.

26. Belyaev, A.O., Karpov, S.A., Keeling, P.J., and Tikhonenkov, D.V. (2024). The nature of ‘jaws’: a new predatory representative of Provora and the ultrastructure of nibbling protists. Open Biol. 14, 12. 10.1098/rsob.240158.

27. Yabuki, A., Eikrem, W., Takishita, K., and Patterson, D.J. (2013). Fine structure of Telonema subtilis Griessmann, 1913: a flagellate with a unique cytoskeletal structure among eukaryotes. Protist 164, 556–569. 10.1016/j.protis.2013.04.004.

28. Brown, M.W., Heiss, A.A., Kamikawa, R., Inagaki, Y., Yabuki, A., Tice, A.K., Shiratori, T., Ishida, K.-H., Hashimoto, T., Simpson, A.G.B., and Roger, A.J. (2018). Phylogenomics places orphan protistan lineages in a novel eukaryotic super-group. Genome Biol. Evol. 10, 427–433. 10.1093/gbe/evy014

29. Schneider, C.A., Rasband, W.S., and Eliceiri, K.W. (2012). NIH Image to ImageJ: 25 years of image analysis. Nat. Methods 9, 671–675. 10.1038/nmeth.2089.

30. Picelli, S., Faridani, O.R., Björklund, A.K., Winberg, G., Sagasser, S., and Sandberg, R. (2014). Full-length RNA-seq from single cells using Smart-seq2. Nat. Protoc. 9, 171– 181. 10.1038/nprot.2014.006.

31. Kolisko, M., Boscaro, V., Burki, F., Lynn, D.H., and Keeling, P.J. (2014) Single-cell transcriptomics for microbial eukaryotes. Curr. Biol. 24, R1081–1082. 10.1016/j.cub.2014.10.026.

32. Song, L., and Florea, L. (2015). Rcorrector: efficient and accurate error correction for Illumina RNA-seq reads. GigaScience 4, 48. 10.1186/s13742-015-0089-y.

33. Bolger, A.M., Lohse, M., and Usadel, B. (2014). Trimmomatic: a flexible trimmer for Illumina sequence data. Bioinformatics 30: 2114–2120. B10.1093/bioinformatics/btu170.

34. Bushmanova, E., Antipov, D., Lapidus, A., and Prjibelski, A.D. (2019). rnaSPAdes: a de novo transcriptome assembler and its application to RNA-Seq data. GigaScience 8, giz100. 10.1093/gigascience/giz100.

35. Manni, M., Berkeley, M.R., Seppey, M., and Zdobnov, E.M. (2021). BUSCO: assessing genomic data quality and beyond. Curr. Protoc. 1, e323. 10.1002/cpz1.323.

36. Haas, B.J., Papanicolaou, A., Yassour, M., Grabherr, M., Blood, P.D., Bowden, J., Couger, M.B., Eccles, D., et al. (2013). De novo transcript sequence reconstruction from RNA-seq using the Trinity platform for reference generation and analysis. Nat. Protoc. 8, 1494–1512. 10.1038/nprot.2013.084.

37. Lang, B.F., Beck, N., Prince, S., Sarrasin, M., Rioux, P., and Burger, G. (2023). Mitochondrial genome annotation with MFannot: a critical analysis of gene identification and gene model prediction. Front. Plant Sci. 14, 1222186. 10.3389/fpls.2023.1222186.

38. Seemann, T. (2014). Prokka: rapid prokaryotic genome annotation. Bioinformatics 15, 2068–2069. 10.1093/bioinformatics/btu153.

39. Greiner, S., Lehwark, P., and Bock, R. (2019). OrganellarGenomeDRAW (OGDRAW) version 1.3.1: expanded toolkit for the graphical visualization of organellar genomes. Nucleic Acids Res. 46, W59–W64. 10.1093/nar/gkz238.

40. Guillou, L., Bachar, D., Audic, S., Bass, D., Berney, C., Bittner, L., Boutte, C., Burgaud, G., de Vargas, C., Decelle, J., et al. (2013). The Protist Ribosomal Reference database (PR2): a catalog of unicellular eukaryote Small Sub-Unit rRNA sequences with curated taxonomy. Nucleic Acids Res. 41, D597–D604. 10.1093/nar/gks1160.

41. Edgar, R.C. (2010). Search and clustering orders of magnitude faster than BLAST. Bioinformatics 26, 2460–2461. 10.1093/bioinformatics/btq461.

42. Katoh, K., Misawa, K., Kuma, K., and Miyata, T. (2002). MAFFT: a novel method for rapid multiple sequence alignment based on fast Fourier transform. Nucleic Acids Res. 30, 3059–3066. 10.1093/nar/gkf436.

43. Capella-Gutiérrez, S., Silla-Martínez, J.M., and Gabaldón, T. (2009). trimAl: a tool for automated alignment trimming in large-scale phylogenetic analyses. Bioinformatics 25, 1972–1973. 10.1093/bioinformatics/btp348.

44. Kozlov, A.M., Darriba, D., Flouri, T., Morel, B., and Stamatakis, A. (2019). RAxML-NG: a fast, scalable and user-friendly tool for maximum likelihood phylogenetic inference. Bioinformatics 35, 4453–4455. 10.1093/bioinformatics/btz305.

45. Letunic, I., and Bork, P. (2021). Interactive Tree of Life (iTOL) v5: an online tool for phylogenetic tree display and annotation. Nucleic Acids Res. 49, W293–W296. 10.1093/nar/gkab301.

46. Richter, D.J., Berney, C., Strassert, J.F.H., Poh, Y.-P., Herman, E.K., Muñoz-Gómez, S.A., Wideman, J.G., Burki, F., and de Vargas, C. (2022). EukProt: a database of genome-scale predicted proteins across the diversity of eukaryotes. Peer Community J. 2, e56. 10.24072/pcjournal.173.

47. Burki, F., Corradi, N., Sierra, R., Pawlowski, J., Meyer, G.R., Abbott, C.L., and Keeling, P.J. (2013) Phylogenomics of the intracellular parasite Mikrocytos mackini reveals evidence for a mitosome in Rhizaria. Curr. Biol. 23, 1541–1547. 10.1016/j.cub.2013.06.033.

48. Minh, B.Q., Schmidt, H.A., Chernomer, O., Schrempf, D., Woodhams, M.D., con Haeseler, A., and Lanfear, R. (2020). IQ-TREE 2: new models and efficient methods for phylogenetic inference in the genomic era. Mol. Biol. Evol. 37, 1530–1534. 10.1093/molbev/msaa015.

49. Lartillot, N., Lepage, T., and Blanquart, S. (2009). PhyloBayes 3: a Bayesian software package for phylogenetic reconstruction and molecular dating. Bioinformatics 25, 2286– 2288. 10.1093/bioinformatics/btp368.

