## Supplementary figures and images for "Phylogenetic position and mitochondrial genome evolution of ‘orphan’ eukaryotic lineages"

### Supplementary figure 1

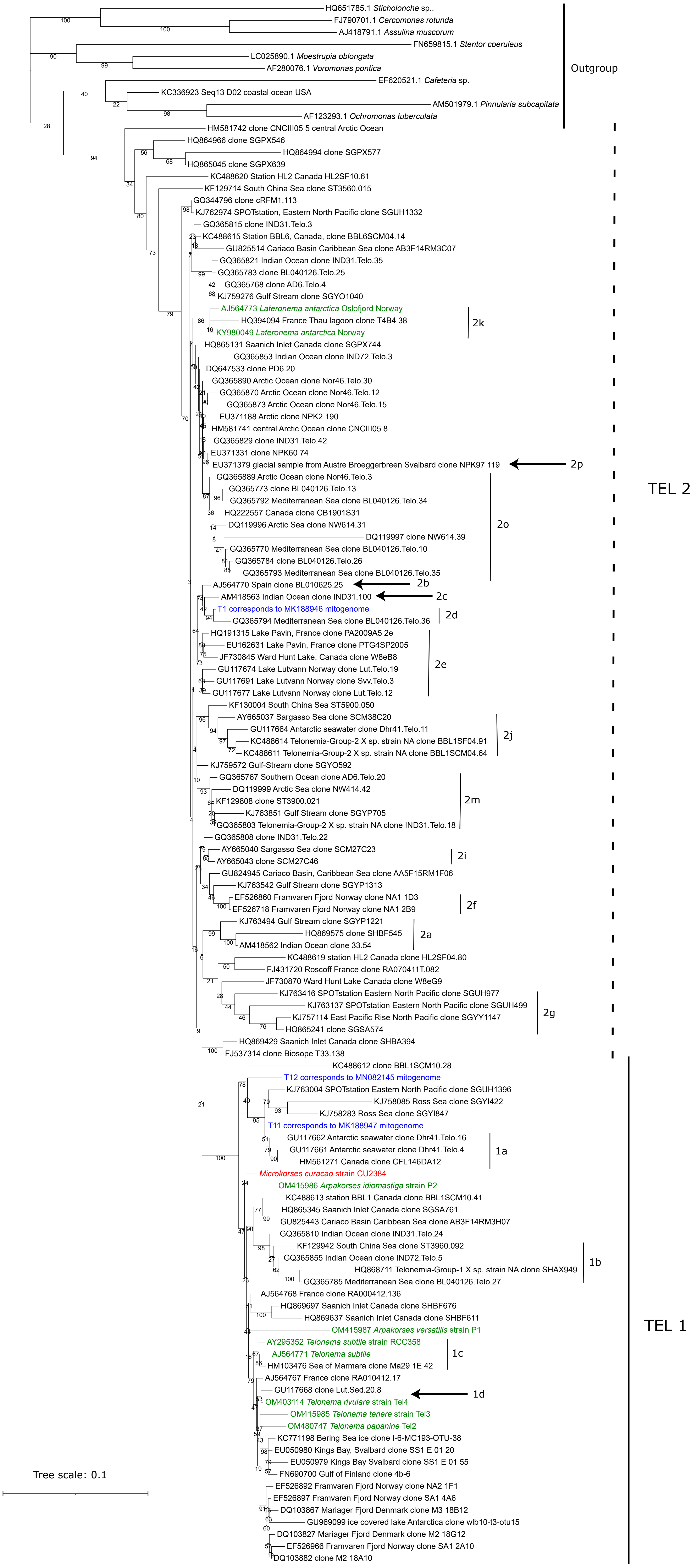

### Supplementary figure 2

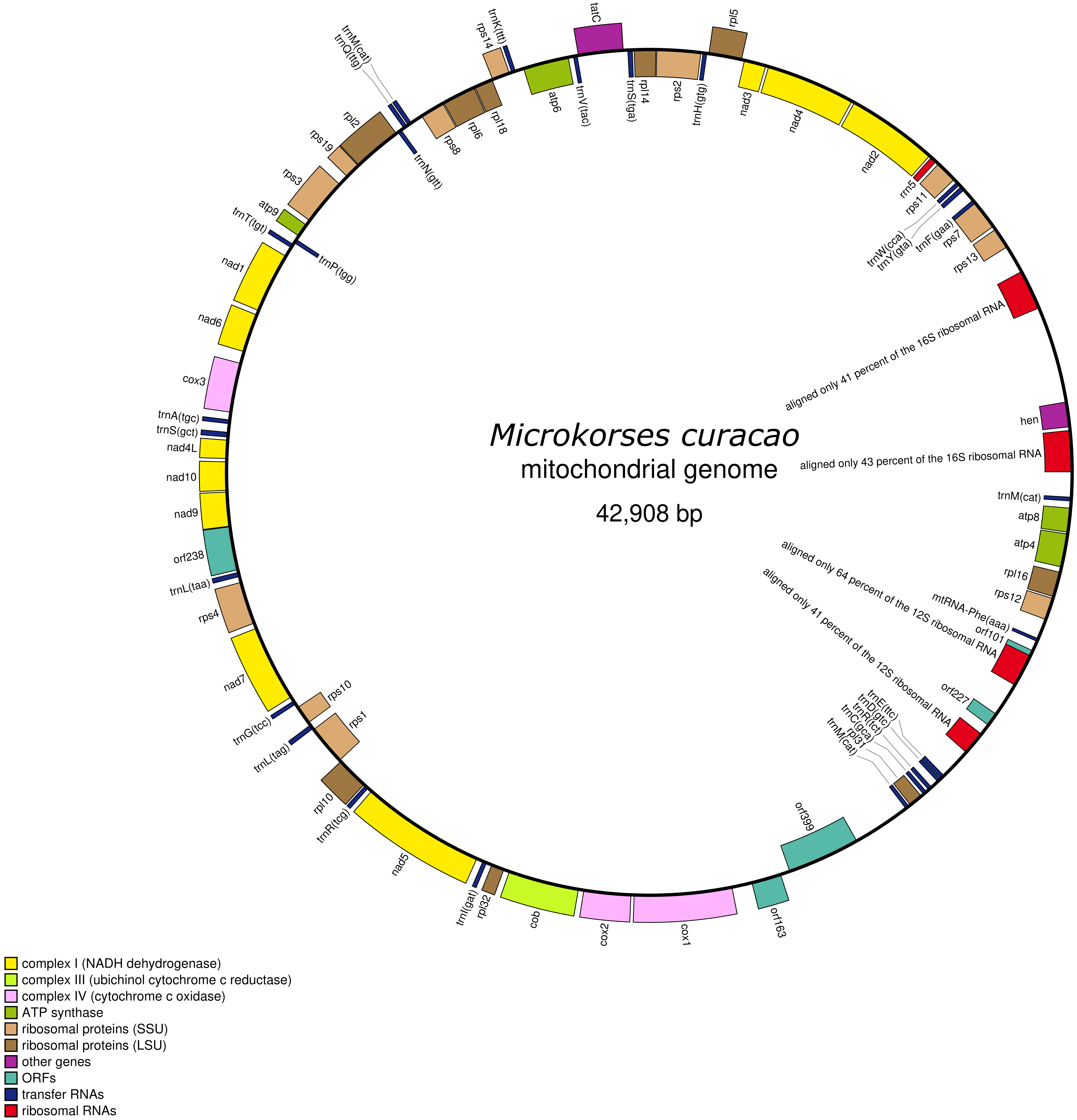

### Supplementary figure 3

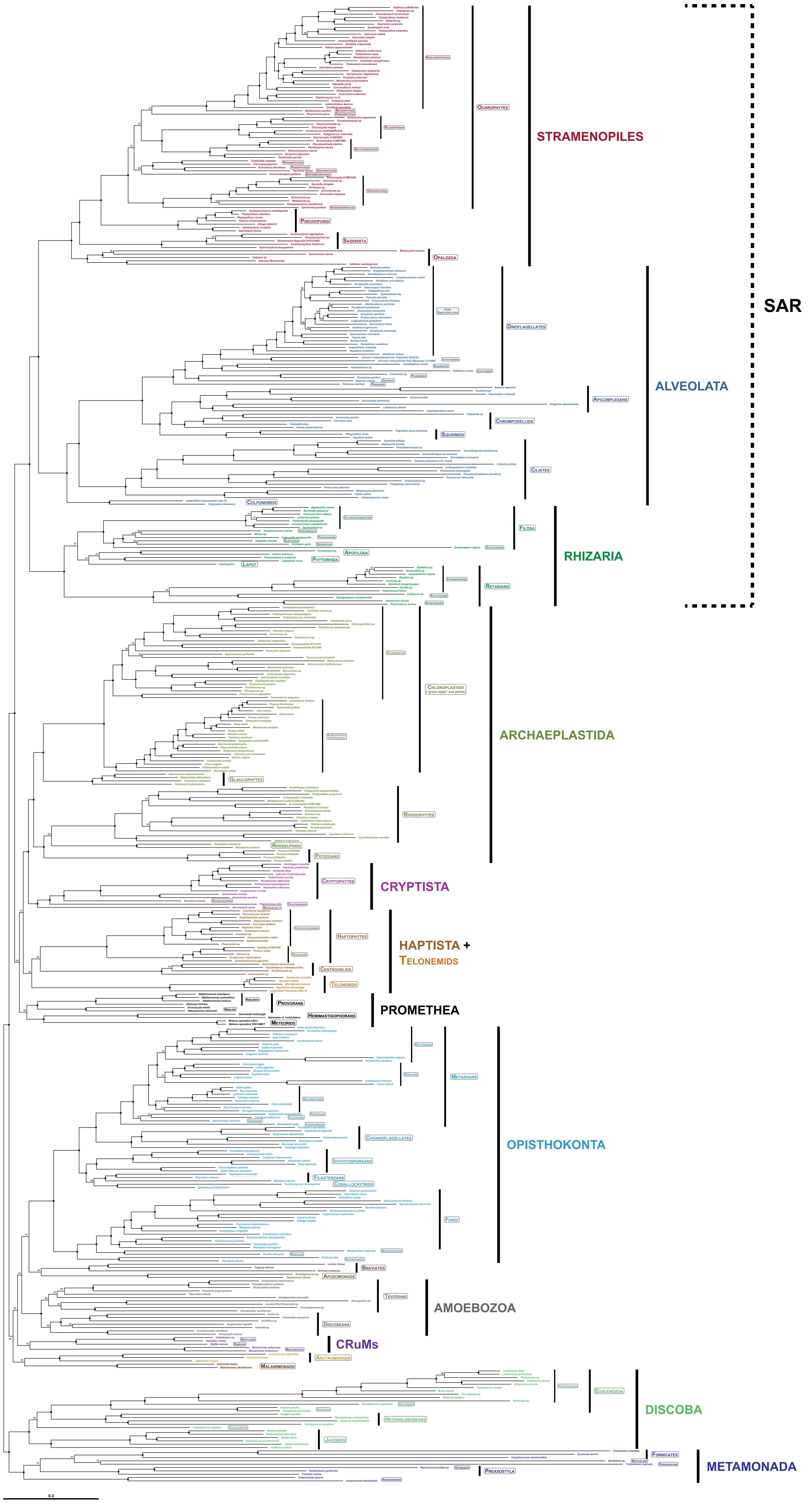

### Supplementary figure 4

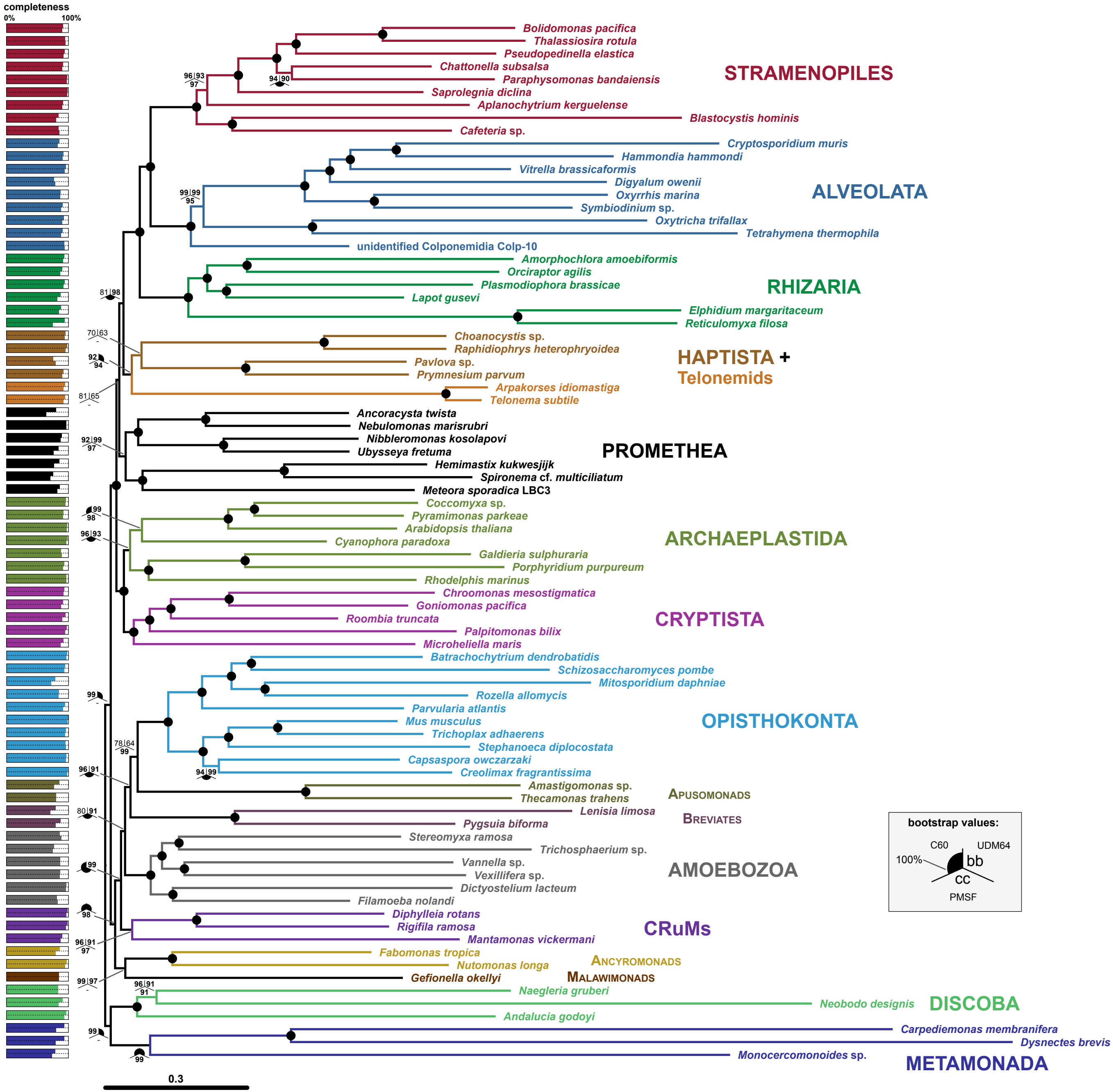

### Supplementary figure 5

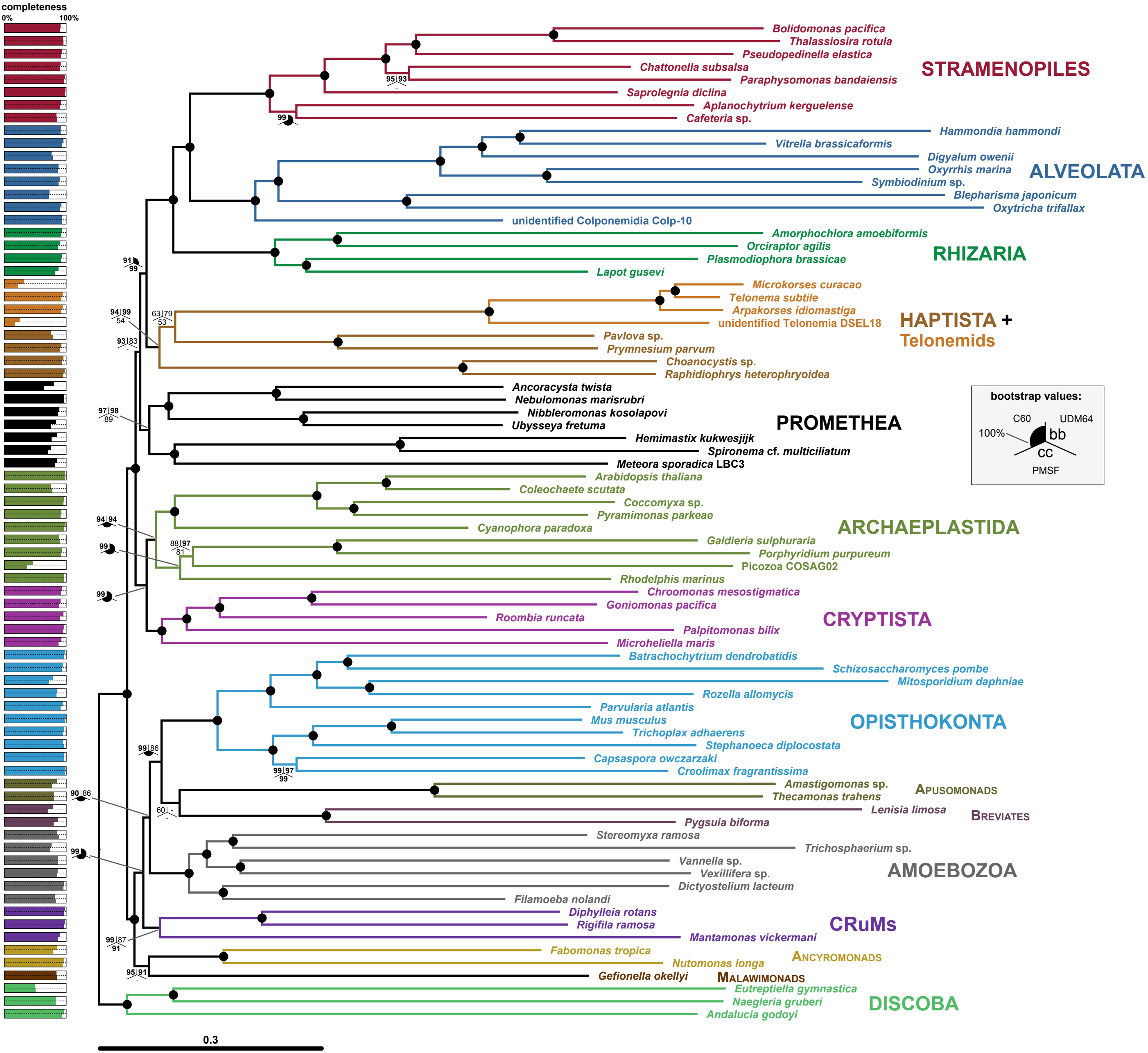

### Supplementary figure 6

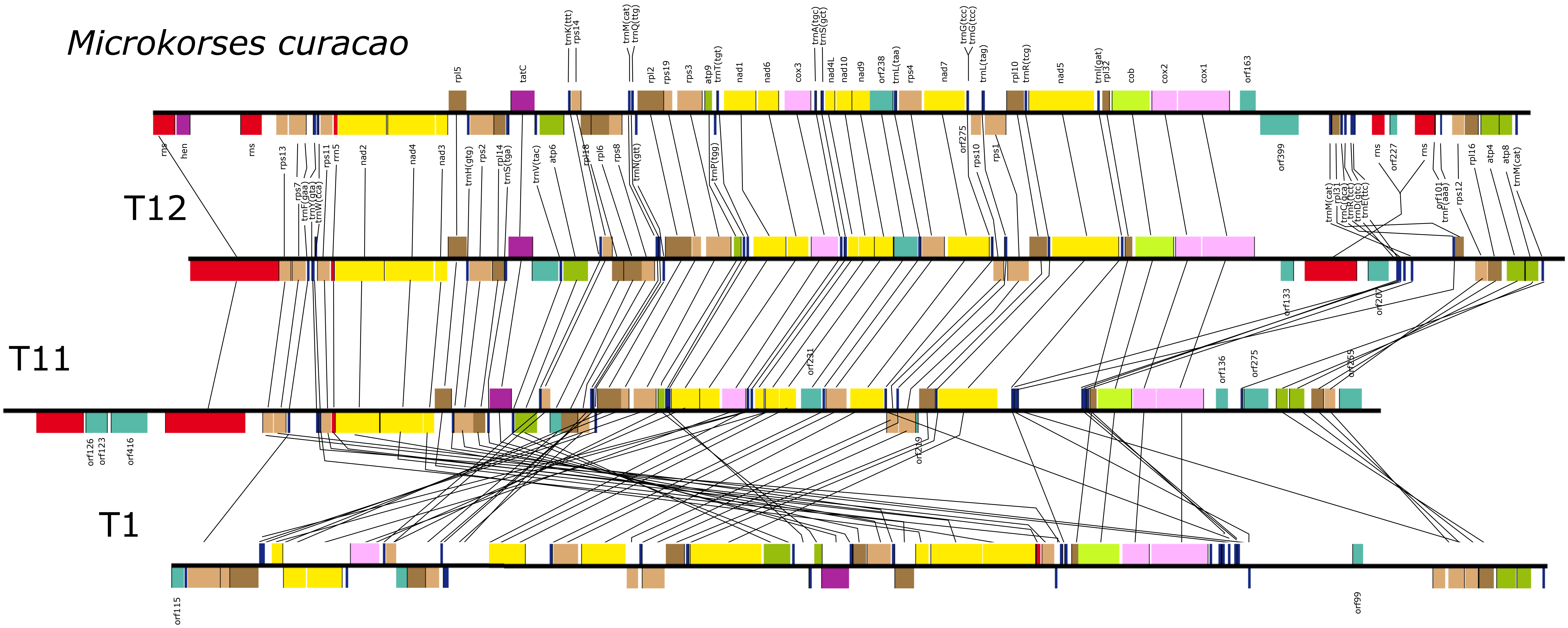
