## Supplementary discussion for "Phylogenetic position and mitochondrial genome evolution of ‘orphan’ eukaryotic lineages"

**Comparison of *Microkorses curacao* gen. et sp. nov. with other described telonemids**

Morphological descriptions are currently available for seven telonemid species, namely *Arpakorses versatilis*, *Arpakorses idiomastiga*, *Telonema papanine*, *Telonema tenere*, *Telonema rivulare*, *Telonema subtile*, and *Lateronema antarctica*. The set of morphological characters observed in *Microkorses curacao* gen. et sp. nov. is a mosaic and doesn’t clearly fit the taxonomic framework, probably due to a considerable number of yet unstudied telonemid representatives, as evidenced by the diverse array of the environmental SSU rRNA gene sequences available (Suppl. Fig. 1). *Telonema subtile*^8^ (original spelling *subtilis*) was for a long time the only described species of the group. Its main characters included a tear-shaped cell body with equal posteriorly directed flagella at the pointed end; it was reportedly 6-8 µm long, with a maximal width of 3-4 µm. The length of the flagella was about the same as that of the cell body. Long after, a second species, *Telonema antarcticum*, was described^50^, but a decade later it was separated as the sole member of genus *Lateronema*^51^. The length (8-16 µm) and width (6-12 µm) of its pyriform cell were notably bigger compared to *T. subtile,* the unequal flagella considerably exceeded the cell body in length, and one flagellum was directed laterally during the swimming. Justifications for establishing a separate genus included the different length of flagellar acronemes, flagella position during swimming, presence of alveoli, absence of isodiametric extrusomes, and presence of tripartite flagellar hairs^51^. Subsequently, Tikhonenkov et al.^16^ provided a morphological description of three new *Telonema* species and two species in the new genus *Arpakorses.* The morphological identity of *Telonema*, as a result, was significantly blurred, but not redefined by the authors. The newly described species were bigger compared to *T. subtile* (*T. papanine*: 8.5–13.7 ✕ 4.2–8.3 µm; *T. tenere*: 5.8–10.2 ✕ 3.5–7.2 µm; *T. rivulare*: 10–14 ✕ 5.0–9.5 µm), had flagella shorter than the cell body, and two of them (*T. papanine* and *T. rivulare*) were isolated from freshwater as opposed to the marine habitat of all previously described telonemids. This left only the equal length of the flagella as a unifying character of the genus and made its definition mostly phylogeny-based (all four species form a clade, which includes multiple environmental sequences). Considerations of *Arpakorses* make the picture even more complex, since species in this genus are similar in size to *T. subtile,* have flagella either longer (*A. versatilis*) or shorter (*A. idiomastiga*) than the cell body, have either equal flagella as in *Telonema* (*A. idiomastiga*) or unequal flagella as in *Lateronema* (*A. versatilis*); either possess tripartite mastigonemes like *Lateronema* (*A. versatilis*) or don’t (*A. idiomastiga*). On top of this, the two species do not group together in the published SSU rRNA gene tree (see Fig. 1 in Tikhonenkov et al.^16^), nor do they in our analysis (Suppl. Fig. 1). *Microkorses curacao* described here stands out by its small size (5–6 ✕ 2.5–3 µm) – smaller than all previously described telonemids. Similarly to *Telonema* spp. and *A. idiomastiga*, it has flagella of equal length, notably exceeding the cell length as in *Lateronema* and *A. versatilis*; similarly to *Arpakorses* spp., the cells were mostly substratum-associated and were solitary (unlike *A. idiomastiga*, but similarly to *A. versatilis*). Finally, on the SSU rRNA gene tree *M. curacao* grouped with *A. idiomastiga* and far from the type species of *Arpakorses* (*A. versatilis*), suggesting that *Arpakorses* is not only morphologically heterogenous, but also non monophyletic. Thus, we decided to describe our strain as a member of the separate new genus *Microkorses.* Whether *A. idiomastiga* should be transferred to *Microkorses* or deserves its own generic name should be defined by further analysis, since the grouping of *M. curacao* and *A. idiomastiga* we revealed is weakly supported (Suppl. Fig. 1).

Zoobank lsid of the publication:

urn:lsid:zoobank.org:pub:0D2A9438-3613-47C5-8C8A-ACFFDFB45E60.

*Microkorses* gen. nov. Zlatogursky 2025

urn:lsid:zoobank.org:act:BA53FCE3-EA3A-4F32-828C-47BE5F6B1D06

**Etymology:** *Microkorses* originates from Ancient Greek μικρός (mikrós, “small, little, micro-”) and Greek κορσές (corset).

**Type species:** *Microkorses curacao*.

**Diagnosis:** Telonemids with acronematic flagella of equal length exceeding that of cell body. Cells attached or swimming in spiral trajectory, solitary.

*Microkorses curacao* sp. nov. Zlatogursky 2025

urn:lsid:zoobank.org:act:0147F213-46F3-40B3-B333-8A19DFFAC762

**Etymology:** *curacao* - for the type locality.

**Type locality:** plankton of Caribbean Sea near water factory dive site at Willemstad, Curaçao (grid reference 12.108993807580502, -68.95353029793979).

**Diagnosis:** Cell length 5-6 µm, cell width in widest part 2.5-3 µm. Flagella acronematic, of equal length (unlike *Arpakorses versatilis*) when observed with light microscopy. The cells attach to the substratum and remain motionless except for a slight trembling or display circular movement (unlike *Arpakorses idiomastiga*). When swimming, cell body rotating around central axis, changing the direction of central axis rapidly, resulting in spiral trajectory. Cells solitary, not in clusters.

**Type figure:** Fig. 1 of the present publication. See also Supplementary videos S1-3.

**Gene sequence**: The SSU rRNA gene sequence - GenBank accession number to be inserted prior to publication.

**Mitochondrial genome features of telonemids**

The obtained *Microkorses curacao* mitogenome is the first culture-based mitochondrial genome characterised for any telonemid, however three other telonemid mitogenomes were obtained previously, using a culture-free approach: cell sorting and subsequent single-amplified genomes (SAGs) sequencing^19^. The analysis of SSU rRNA gene sequences shows that SAGs T11 and T12 are related to the TEL1 clade (1a subclade), while SAG T1 is attributed to a distantly related TEL2 clade (2d subclade) (Suppl. Fig. 1; clade names are based on Fig. 1 in Tikhonenkov et al.^16^). The overall gene composition of our culture-based and three SAG-based mitogenomes was similar except for the *rpl31* gene missing in T11 and T1 and some differences in transfer RNAs content (Suppl. Fig. S4). In particular, *M. curacao* had a redundant set of two phenylalanine transfer RNAs: *trnF(aaa)* and *trnF(gaa*), while only *trnF(gaa)* was present in T1, T11 and T12. Similarly, all four genomes contained isoleucine transfer RNA *trnI(gat)*, but T1 in addition redundantly had *trnI(aau)*. All the transfer RNA genes in all four genomes were duplicated except *trnM(cat)* which was always present in six copies per genome. The above-mentioned additional *trnF(aaa)* in *M. curacao* was present in single copy, while the redundant *trnI(aau)* in T1 was present in two copies (Suppl. Table S1). The synteny with the bacterial-like genomes of jakobids, noted previously (Wideman et al., 2020) was also present in *Microkorses*. In general, the genome architecture was considerably similar among TEL1 mitogenomes (*Microkorses*, T11, T12), but compared to them it was notably rearranged in TEL2 (represented by T1) (Suppl. Fig. S6).

T11, T12 and *Microkorses* had a similarly long ORF between *rps4* and *nad9*, which was missing in T1, potentially representing a synapomorphy of the TEL1 clade. T12 had a unique ORF, *orf265*, between *atp6* and *tatC*, missing in other mitogenomes. T1’s *orf99* revealed some homology to a bacterial cell-wall-associated hydrolase. Notably, the gene for the RNA of the large ribosomal subunit (*rnl*) in *M. curacao* contained a group I intron, encoding a LAGLIDADG homing endonuclease, which is also known for other protists, notably provorans and centrohelids^5,52^.

**Supplementary references**

50. Klaveness, D., Shalchian-Tabrizi, K., Thomsen, H.A., Eikrem, W., and Jakobsen, K.S. (2005). *Telonema antarcticum* sp. nov., a common marine phagotrophic flagellate. Int. J. Syst. Evol. Microbiol. 55, 2595–2604. 10.1099/ijs.0.63652-0.

51. Cavalier-Smith, T., Chao, E.E., and Lewis, R. (2015). Multiple origins of Heliozoa from flagellate ancestors: new cryptist subphylum Corbihelia, superclass Corbistoma, and monophyly of Haptista, Cryptista, Hacrobia and Chromista. Mol. Phylogenet. Evol. 93, 331–362. 10.1016/j.ympev.2015.07.004.

52. Nishimura, Y., Shiratori, T., Ishida, K.I., Hashimoto, T., Ohkuma, M., and Inagaki, Y. (2019). Horizontally-acquired genetic elements in the mitochondrial genome of a centrohelid *Marophrys* sp. SRT127. Sci. Rep. 9, 4850. 10.1038/s41598-019-41238-6.
